# Myofiber stretch induces tensile and shear deformation of muscle stem cells in their native niche

**DOI:** 10.1101/2020.01.14.902510

**Authors:** Mohammad Haroon, Jenneke Klein-Nulend, Astrid D. Bakker, Jianfeng Jin, Carla Offringa, Fabien Le Grand, Lorenzo Giordani, Karen J. Liu, Robert D. Knight, Richard T. Jaspers

## Abstract

**Background:** Muscle stem cells (MuSCs) are requisite for skeletal muscle regeneration and homeostasis. Proper functioning of MuSCs, including activation, proliferation, and fate decision, is determined by an orchestrated series of events and communication between MuSCs and their niche consisting of the host myofiber and neighbouring cells. A multitude of biochemical stimuli are known to regulate fate and function of MuSCs. However, in addition to biochemical factors, it is conceivable that MuSCs residing between basal lamina and sarcolemma of myofibers are subjected to mechanical forces during muscle stretch-shortening cycles due to myofascial connections between MuSCs and myofibers. MuSCs have been shown to respond to mechanical forces *in vitro* but it remains to be proven whether physical forces are also exerted on MuSCs in their native niche and whether they contribute to the functioning and fate of MuSCs.

**Methods:** MuSCs deformation in their native niche resulting from mechanical loading of *ex vivo* myofiber bundles were visualized utilizing *mT/mG* double-fluorescent Cre-reporter mouse and multiphoton microscopy. MuSCs were subjected to 1 hour pulsating fluid shear stress with a peak shear stress rate of 8.8 Pa/s. After treatment, nitric oxide and mRNA expression levels of genes involved in regulation of MuSC proliferation and differentiation were determined.

**Results:** *Ex vivo* stretching of extensor digitorum longus and soleus myofiber bundles caused compression as well as tensile and shear deformation of MuSCs in their niche. MuSCs responded to pulsating fluid shear stress *in vitro* with increased nitric oxide production and an upward trend in *iNOS* mRNA levels, while *nNOS* expression was unaltered. Pulsating fluid shear stress enhanced gene expression of *c-Fos, Cdk4*, and *IL-6*, while expression of *Wnt1, MyoD, Myog, Wnt5a, COX2, Rspo1, Vangl2, Wnt10b*, and *MGF* remained unchanged.

**Conclusions:** We conclude that MuSCs in their native niche are subjected to force-induced deformations due to myofiber stretch-shortening. Moreover, MuSCs are mechanoresponsive as evident by pulsating fluid shear stress-mediated expression of factors by MuSCs known to promote proliferation.

## BACKGROUND

Skeletal muscle regeneration requires activation and proliferation of muscle stem cells (MuSCs). During organismal aging, the number of MuSCs is reduced and MuSCs lose their proliferation potential [1]. Moreover, muscle pathologies characterized by impaired regeneration by MuSCs lead to atrophy and fibrosis [2]. In young, un-injured muscles, myofiber hypertrophy is accompanied by MuSC activation, proliferation, and subsequent fusion with the host myofiber to increase myonuclei numbers [3]. This ensures that the myonuclear domain does not exceed a critical size [3]. Understanding the mechanisms underlying MuSC function is required for adequate strategies to maintain MuSC self-renewal and regenerative function.

During regeneration a multitude of growth factors and cytokines are secreted by myofibers, neighboring immune cells, and fibroblasts, which activate MuSCs and stimulate proliferation, differentiation, and self-renewal [4]. During strenuous exercise, myofibers are subjected to large stretch-shortening deformations by active contraction of myofibers and passive external loads applied to myofibers from antagonist muscle activity. Such mechanical loading stimulates the expression of growth factors and cytokines within the myofiber micro-environment [5]. These growth factors and cytokines act in a paracrine or endocrine manner on the MuSCs and thus regulate their function [6, 7]. This indicates that mechanical loads are essential for MuSC regenerative function, but little is known about how MuSCs themselves respond to mechanical force *in vivo*.

Skeletal muscle fibers have the ability to sense mechanical cues via transmembrane complexes and stretch-activated calcium (Ca^2+^) channels [5]. Three dimensional (3D)-finite element modeling of muscle revealed that global strain application onto a muscle *in vivo* causes high local tensile strains within the extracellular matrix (ECM) [8]. The contractile apparatus within the myofiber is also exerting force laterally [9], and finite element modeling suggests that these lateral forces cause shear deformations of the ECM by slippage along a plane parallel to the imposed stress [10]. These deformations cause interstitial fluid movement within the muscle [11], and currently it is unknown whether MuSCs can sense these fluid movements. Moreover, force induced ECM deformations could also be transmitted to the MuSCs, which are anchored to the sarcolemma of the myofiber by cadherins, and on their apical side to the basal lamina of the endomysium via glycocalyx, integrins, syndecans, dystroglycans, and sarcoglycans [12]. Therefore, it has been suggested that MuSCs experience tensile strain and shear stress following muscle tissue contraction [13]. *In vitro* studies have shown that MuSCs are sensitive to tensile forces [14, 15], however it is unknown whether MuSCs within their native niche are subjected to mechanical forces resulting in cell deformation. In this study, we tested whether physical load induces deformation of MuSCs in their native niche using isolated mouse myofiber bundles containing an intact endomysium and basal lamina. We were able to show that upon myofiber bundle stretch, the MuSCs undergo substantial shear and tensile strain deformation.

Since MuSCs in their niche are mechanically loaded which causes interstitial fluid movement within the muscle [11], the question arises whether they are able to respond to fluid shear-induced mechanical loads by changing the expression of factors involved in MuSC self-renewal, proliferation, and differentiation. Bone cells are subjected to shear forces due to the interstitial fluid flow within the canaliculi of bones during mechanical loading which are estimated to be 0.8-3 Pa [16]. Bone cells respond to these shear forces by prostaglandin production and NO upregulation. Mechanical loading of myofibers has been shown to increase nitric oxide (NO) production and regulate NO synthase (NOS) expression and activity [17]. Pulsating fluid shear stress (PFSS) increases NO production and expression of interleukin-6 (*IL-6*) and cyclooxygenase-2 (*COX2*) in myotubes [18]. NO plays a role in Vang-like protein 2 (Vangl2) expression in MuSCs resulting in enhanced MuSC self-renewal during skeletal muscle regeneration [19]. Non-canonical Wnt signaling plays a crucial role in muscle regeneration [20], and mechanical loading is known to activate Wnt signaling in bones [21]. Mechanical cues, such as tensile strain and shear stress, constitute a physiologically relevant and novel mechanism that can therefore potentially activate Wnt/planar cell polarity (PCP) signaling, thereby promoting MuSC self-renewal. Whether mechanical forces do induce strain and stress onto MuSCs in muscle has yet to be explored. We therefore investigated whether mechanical loads on MuSCs induce NO production and expression of biochemical factors that are crucial for MuSC regenerative function.

## METHODS

### Transgenic mice and myofiber bundle isolation

Animal procedures were conducted according to UK Home Office project license P8D5E2773 (KJL). Paired box 7 (Pax7) creERT2 mice (Jackson strain: Pax7^tm2.1(cre/ERT2)Fan^) on a mixed C57BL6/CD1 background were crossed with *mTmG* reporter mice (Jackson strain: Gt(ROSA)26Sor^tm4(ACTB-tdTomato,-EGFP)Luo^). Targeted deletion in the CreERT animals was performed by intraperitoneal tamoxifen injection (150 mg/kg body weight) at 40 and 20 h prior to euthanasia. Twenty mg/ml tamoxifen solution was prepared by dissolving 10 mg tamoxifen in 100 µl 100% ethanol and adding 900 µl corn oil. Compound heterozygous transgenic mice (15-17 months, n=3) were used for *in situ* mechanical loading of MuSCs and image acquisition as described below. Mice were sacrificed, and extensor digitorum longus (EDL) and soleus (SO) muscles extracted. Both EDL and SO muscles were dissected for isolation of myofiber bundles, consisting of ∼20 myofibers, using fine-tipped forceps and scissors under a dark field illuminated microscope.

### Primary MuSC isolation and fluorescence-activated cell sorting

Primary MuSCs were isolated from young (2 months; n=4) male mouse (C57BL/6J) hindlimb muscles (Hamstring muscle group, Quadriceps, Tibialis, Extensor digitorum longus, Gastrocnemius, Soleus, Gluteus) after enzymatic digestion followed by Fluorescence-activated cell sorting (FACS) purification. Briefly, after dissection muscles were first digested with collagenase II (1000 U/ml; Worthington Biochemical Corporation, Lakewood, NJ, USA) in Ham’s F10 containing 10% horse serum, washed and then further digested with collagenase II (1000 U/ml) and dispase (11 U/ml) for 30 min. After digestion cells were washed in Ham’s F10 containing 10% horse serum, passed 10 times through a 20-gauge needle syringe and then filtered with a 35-mm cell strainer (Falcon^®^, Corning, NY, USA). Cells were then stained with the following antibodies: rat CD31-eFluor450 (1/500; eBiosciences™, Thermo-Fisher Scientific, San Diego, CA, USA), rat CD45-eFluor450 (1/500; eBiosciences™, Thermo-Fisher Scientific), rat Ly6A-FITC (SCA1) (1/500; eBiosciences™, Thermo-Fisher Scientific), rat CD106-PE (1/200; eBiosciences™, Thermo-Fisher Scientific) and rat α7 integrin-APC (1/1000; AbLab, Vancouver, Canada) and sorted using a FACS Aria II (BD Biosciences, San Jose, CA, USA). Satellite cells were isolated as CD31^−^, CD45^−^, Sca1^−^, α7 integrin^+^, CD106^+^.

### *In situ* mechanical loading of muscle stem cells

The myofiber bundles (n=8) were mounted at their slack length (low strain) by attaching the tendons at both ends to small rods in chambers (Bioptechs, Butler, PA, USA; see Fig. S1) containing tyrode solution (NaCl 7.5 g/l; KCl 0.35 g/l; MgCl_2_.6H_2_O 0.214 g/l; NaH_2_PO_4_.H_2_O 0.058 g/l; NaHCO_2_ 1.7 g/l; CaCl_2_.H_2_O 0.2 g/l) which was saturated with a carbogen gas mixture (95% O_2_ and 5% CO_2_) at 37°C. To induce a high strain on the myofiber bundle, the bundle was stretched by ∼40% above slack length (i.e. high strain) in accordance with the previous literature [22].

### *In situ* image acquisition

Images were acquired on a Zeiss Axioscope (Oberkochen, Germany) using a 7MP multiphoton system with Vision II and MPX (Chameleon, Santa Clara, CA, USA) lasers and a 20x water dipping objective (NA 1.0). The Vision II laser was tuned to 890 nm (range: 880-900 nm), and the MPX laser to 1200 nm with reflected emissions captured by non-descanned photomultiplier detectors (channel 1 and 2), GaAsP BiG detector (channel 3), or transmitted light detector (channel 4). Emission filters used were shortpass 485 nm (channels 1 and 4), bandpass 500-550 (channel 2), and bandpass 575-610 (channel 3). The signals detected were second harmonic (channel 1), GFP (channel 2), and tdTomato (channel 3). Image stacks were acquired every ∼1 µm in Z direction.

### Image analysis

Cells were segmented in the Medical Imaging Interaction Toolkit (MITK, www.mitk.org) for 3D reconstruction. Analysis of cellular parameters, i.e. sarcomere length, cell length, cell width, cell aspect-ratio (major axis/minor axis), and cell roundness (4×([area])/(π×[major axis]^2^) were performed in ImageJ, version 1.52h (Wayne Rasband, National Institutes of Health, Bethesda, MD, USA). Tensile strain was calculated by the change in length (ΔL) divided by the initial length (L_0_) in XY plane (ΔL/L_0_). Shear strain was quantified by measuring the difference in change in length (ΔL) of the two lateral sides of the cell, and the change in angle between both lateral sides in the YZ plane (shear angle).

### Cell culture

For *in vitro* mechanical loading by PFSS, MuSCs were expanded on matrigel (Corning, Bedford, MA, USA)-coated culture flasks with growth medium consisting of Ham’s F-10 Nutrient Mix (Gibco, Paisley, UK) supplemented with 20% fetal bovine serum (FBS, Gibco), 10 µg/ml penicillin (Sigma-Aldrich, St. Louis, MO, USA), 10 µg/ml streptomycin (Sigma-Aldrich), and 2.5 ng/µl of recombinant human fibroblast growth factor (R&D systems, Minneapolis, MN, USA), and cultured at 37°C in a humidified atmosphere of 5% CO_2_ in air. Upon 70% confluence, cells were harvested using 0.1% trypsin and 0.1% EDTA (Gibco) in phosphate-buffered saline.

### Pulsating fluid shear stress

MuSCs were seeded at 12-20×10^3^/cm^2^ on matrigel (Corning)-coated glass slides (2.5×6.5 cm; n =10), and cultured for 2-4 days. One hour before PFSS, culture medium was refreshed by medium containing a low serum concentration (2% FBS). PFSS was applied as described earlier [18]. Briefly, PFSS was generated by pumping 7 ml of culture medium through a parallel-plate flow chamber containing MuSCs. Cells were subjected to a cyclic changing pressure gradient with a peak shear stress rate of 6.5 Pa/s (pulse amplitude: 1 Pa; pulse frequency: 1 Hz). We adapted the 1Hz frequency which mimics the walking strain cycles (stride frequency) [23]. Static control cells were kept in similar conditions as PFSS loaded cells. After 1 h PFSS treatment or static control culture, images were taken and RNA was isolated. To determine the number of MuSCs detached as a result of PFSS exposure, cells were counted in random images acquired pre and post-PFSS.

### Nitric oxide analysis

Medium samples were taken from PFSS-treated and static cell cultures (n = 20) at intervals (10, 30, and 60 min) for NO analysis. NO production was measured as nitrite (NO_2_ ^−^) accumulation in the medium using Griess reagent, and absorbance was measured at 540 nm with a microplate reader (BioRad Laboratories Inc., Veenendaal, The Netherlands) as described earlier [18].

### RNA isolation and reverse transcription

After PFSS and static treatment (n=7-10), micrographs were taken, cells were lysed with 700 µl Trizol (Thermo-Fisher Scientific, Carlsbad, CA, USA), and stored at −80°C overnight. Total RNA was isolated using RiboPure™Kit (Applied Biosystems, Foster City, CA, USA) and quantified (NanoDrop Technologies, Thermo-Fisher Scientific, Wilmington, DE, USA). mRNA (400 ng) was reverse-transcribed to complementary DNA (cDNA) using a High Capacity RNA-to-cDNA kit (Applied Biosystems).

### Quantitative Real-Time PCR

Real-Time PCR was performed on the StepOne™Real-Time PCR system (Applied Biosystems). Primers were designed using Universal Probe library from Roche Diagnostics. Data were analyzed using StepOne™v2.0 software (Applied Biosystems) and normalized relative to *18S* ribosomal RNA levels. *Wnt1, Wnt3a, Wnt5a, Wnt7a, Wnt10b, IL-6, Rspo1, Dkk1, COX2, nNOS, eNOS*, and *iNOS* transcript levels were measured using Taqman^®^ qPCR using inventoried Taqman^®^ gene expression assays (Applied Biosystems). mRNA levels of *18S, MyoD, Myog, c-Fos, Cdk4* (Cyclin-dependent kinase 4), *Vangl2*, and *MGF* were measured using SYBR^®^ green (Thermo-Fisher Scientific). Primer sequences used for real-time PCR: *18S* (5’ GTAACCCGTTGAACCCCATT and 5’ CCATCCAATCGGTAGTAGCG); *MyoD* (5’CATCCAGCCCGCTCCAAC and 5’ GGGCCGCTGTAATCCATCATGCC); *Myog* (5’ CCAGCCCATGGTGCCCAGTGA and 5’ CCAGTGCATTGCCCCACTCCG); *c-Fos* (5’ TCACCCTGCCCCTTCTCA and 5’ CTGATGCTCTTGACTGGCTCC); *Cdk4* (5’ GGGGAAAATCTTTGATCTCATTGGA and 5’ AAGGCTCCTCGAGGTAGAGATA); *Vangl2* (5’ CCCCAGTTCACACTCAAGGT and 5’ ACTTGGGCAGGTTGAGGAG); *MGF* (5’ GGAGAAGGAAAGGAAGTACATTTG and 5’ CCTGCTCCGTGGGAGGCT).

### Statistical analysis

The paired sample t-test was used to test statistical significant differences in MuSC deformation data, the independent t-test to test differences in tensile strain data, and the one-sample t-test to test differences in shear strain data. Differences in NO production data were tested using ANOVA. One-sample t-test was performed for statistical analysis of gene expression data. Data were expressed as mean ± SEM, and p<0.05 was considered significant.

## RESULTS

### *Ex vivo* myofiber stretch induced MuSC deformation

We characterized MuSCs in their niche, within an isolated myofiber bundle from the EDL and SO muscle (Fig. 1). Sarcomere length was ∼2.5 µm, slightly above the slack length of sarcomeres as described earlier (Fig. 2A) [24]. Following stretching of EDL myofibers up to ∼40% above their slack length, MuSCs on the host myofiber increased in length by the same magnitude (Fig. 1A-D; Fig. 2C). Similar increases in sarcomere length and MuSC were observed for myofiber bundles from SO muscles (Fig. 1I-L). MuSCs on EDL myofibers displayed clear cell protrusions, whereas those on SO myofibers did not. Mean tensile strain on MuSCs of stretched EDL and SO myofibers was 43%, which was of the same order of magnitude as the increase in strain of the myofibers (∼40%; Fig. 2B). In addition to lengthening of MuSCs, filopodia were reoriented in the direction of the stretched myofibers as indicated by the change in position of the processes when the myofibers were stretched compared to slack length (Fig. 1E,F). MuSCs of SO myofibers showed less distinct processes compared to MuSCs of EDL myofibers (Fig. 1M,N). A comparison of the displacement of the lateral sides of the MuSCs in the YZ plane, and changes in the angles between the two lateral sides showed that the MuSCs were also subjected to shear deformation (Fig. 1G,H,O,P). The displacement of the lateral sides of MuSCs differed by ∼40%, and the shear angle showed a 12 degrees-displacement from a fixed axis in both EDL and SO myofibers (Fig. 2D,E).

**Figure 1.**
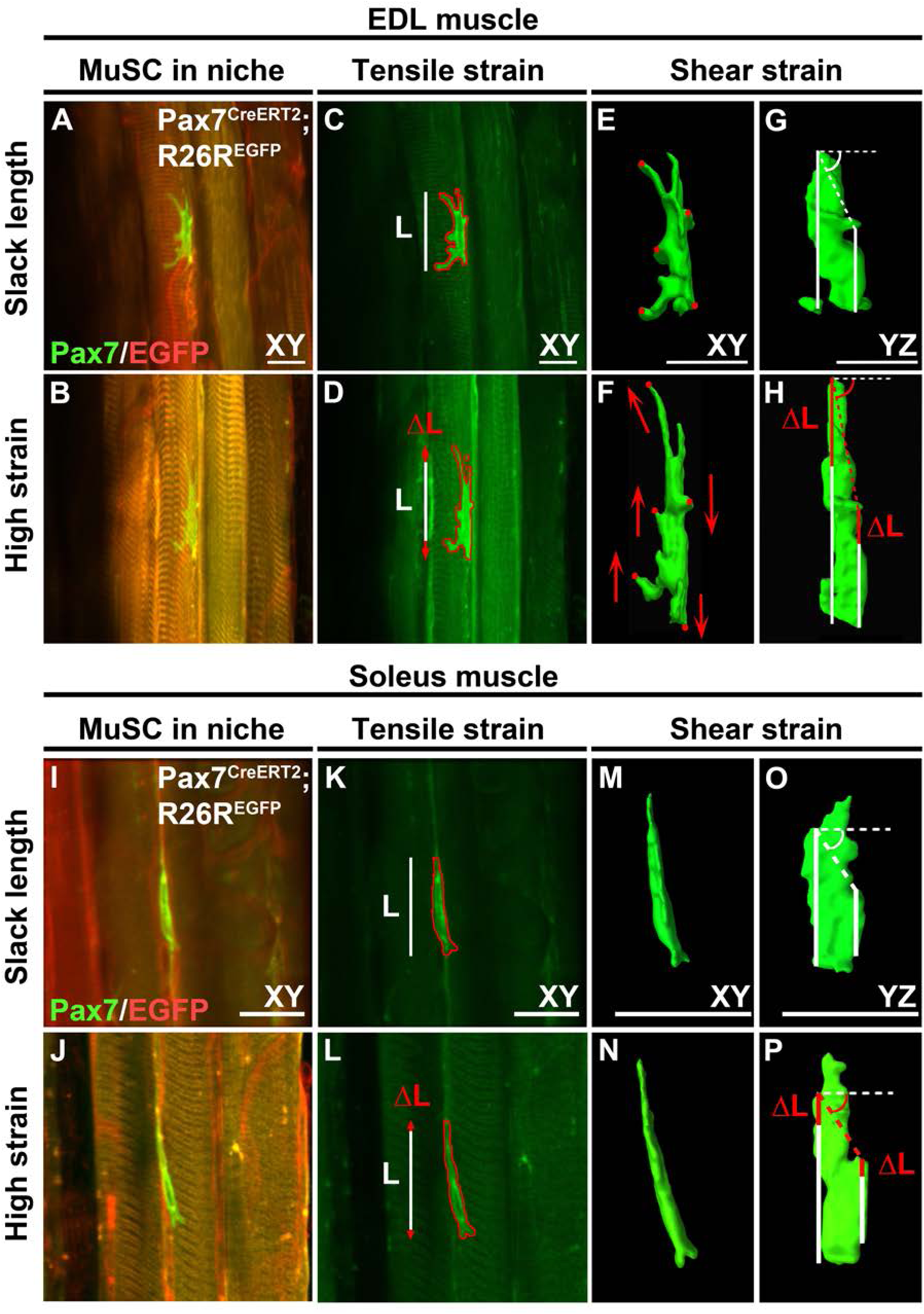
Live cell imaging of MuSCs within their niche at slack length and high strain myofiber bundles. (A, B, I, J) Micrographs of typical MuSCs in their native niche within myofiber bundle at slack length and at high strain (∼40% above slack length) from extensor digitorum longus (EDL) soleus muscles. (C, D, K, L) Measurement of MuSC length change in the XY plane. White lines: length of MuSCs at myofiber bundle slack length. Red arrows: increase in length at extended myofiber bundle length. (E-H, M-P) 3D-reconstructions of MuSCs at slack length and high myofiber strain illustrating MuSC shear deformations in XY and YZ planes. Stretching of the myofiber bundles caused displacement of cell regions (red circles) in the direction indicated by the arrows (red) in the XY plane implicating shearing of MuSC (E, F). (G, H, O, P) Differences in MuSC length changes at both lateral sides indicate an increase in shear angle (i.e. angle of the dashed lines in YZ plane). Scale bar, 20 μm.

**Figure 2.**
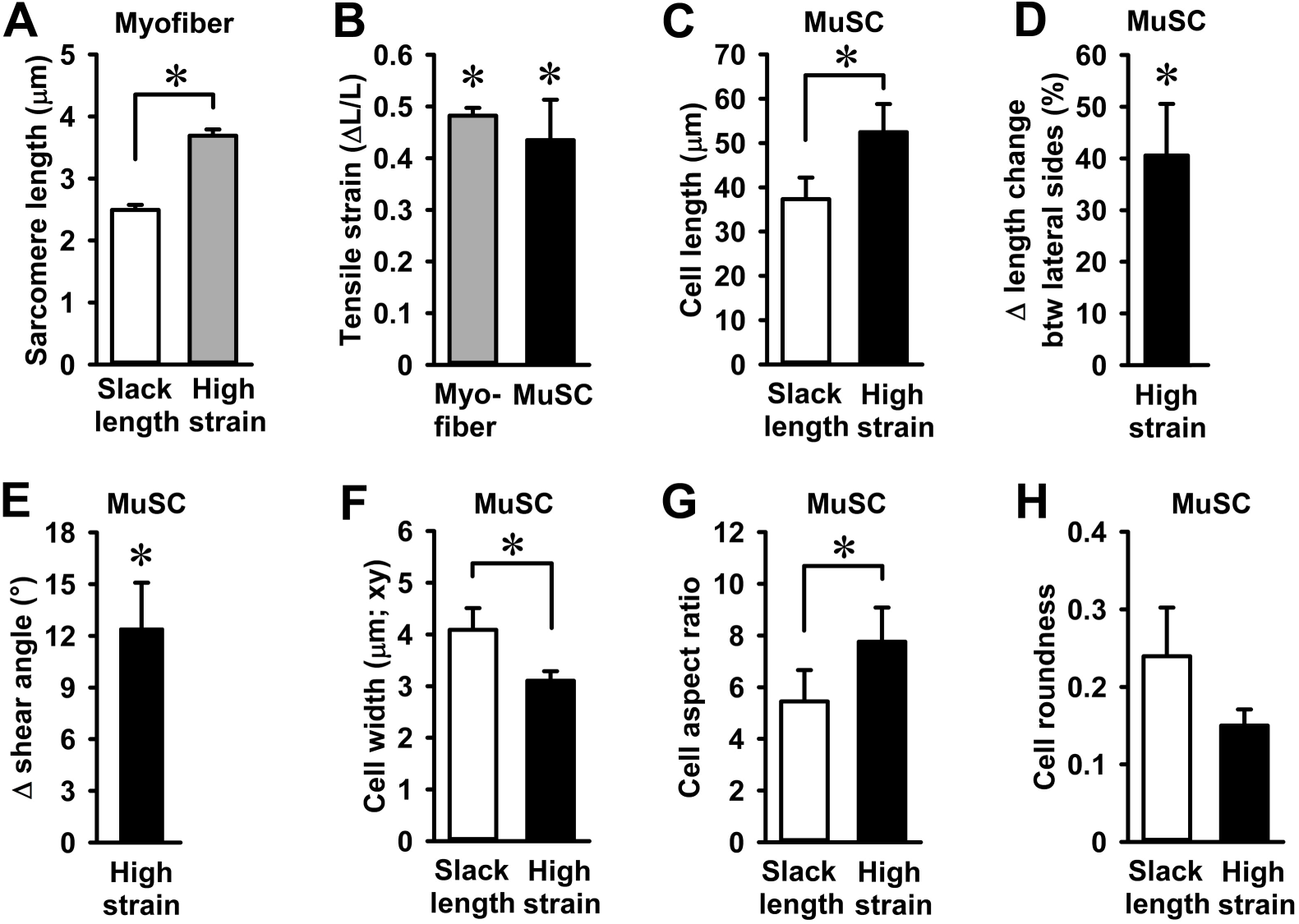
Quantification of myofiber stretch-induced compression, strain, and shearing of MuSCs. (A) Increase in sarcomere length upon EDL and SO myofiber bundle stretch. (B) Myofiber bundle sarcomere strain and MuSC strain were similar. (C) MuSC length at myofiber bundle slack length and high strain. (D) MuSCs experienced shear strain upon myofiber bundle stretch. (E) At high strain, the shear angle between the two lateral sides of the cell also changed by 12 degrees. (F-H) Myofiber stretch decreased cell width and increased MuSC aspect ratio, but did not affect MuSC roundness. n=6-8 cells. Btw; between. *Significant effect of stretch, p<0.05.

Stretching of both EDL and SO myofiber bundles induced MuSC compression and decreased MuSC width by ∼21% (Fig. 2F). Changes in cell morphology after stretch indicated an increase in MuSC aspect-ratio, as MuSCs became more elongated while a negative trend in roundness was noted (Fig. 2G,H). These results showed that MuSCs within their native niche were subjected to tensile and shear forces causing corresponding deformations.

### MuSCs were mechanosensitive to pulsating fluid shear stress

Mechanical force-induced deformation of cell shape may induce specific molecular responses by MuSCs, including elevated NO production. We first determined whether PFSS caused detachment of MuSCs from the substratum. Micrographs after static and PFSS treatment indicated that ∼15% of MuSCs detached after PFSS treatment, whereas no difference was observed in the static controls (Fig. 3A). As NO is produced by NOS, we assessed whether MuSCs express the different isoforms of NOS (*iNOS, eNOS, nNOS*). We found that myoblasts express *nNOS* and *iNOS*, but *eNOS* transcripts were not detectable (Fig. 3B). PFSS did not affect the expression of *nNOS*, but an upward trend was seen for *iNOS* mRNA levels after PFSS treatment (Fig. 3B). To assess NO production by MuSCs, NO levels in the culture medium were quantified. PFSS resulted in increased NO production by MuSCs (2-2.5 fold) after 10, 30, and 60 min (Fig. 3C). These results indicated that MuSCs were sensitive to PFSS, and that NO production was most likely related to alterations in iNOS enzyme activity.

**Figure 3.**
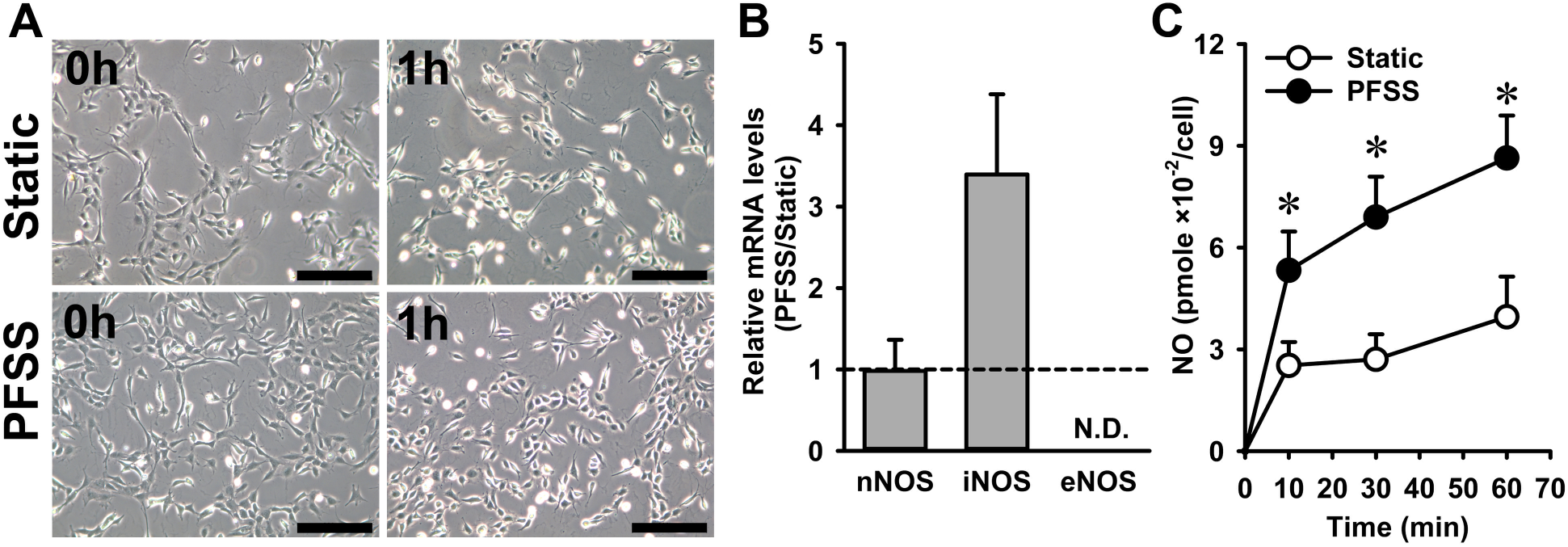
Pulsating fluid shear stress induced NO production and elevated *iNOS* expression in primary MuSCs. (A) Micrographs of cultured primary mouse MuSCs at 1 h post static and PFSS treatment indicate ∼15% loss of cells due to PFSS application. (B) PFSS did not change the expression of *nNOS*, while *eNOS* was not detectable, and PFSS caused an upward trend in the level of *iNOS* mRNA transcripts. (C) PFSS induced NO production at 10, 30, and 60 min by 2-2.5 fold. n=20. PFSS, pulsating fluid shear stress; ND, not detectable. *Significant effect of PFSS, p<0.05. Scale bar, 200 μm.

### PFSS upregulated *c-Fos* and *Cdk4* with no effect on *MyoD* and myogenin (*Myog*)

To assess whether PFSS induces alterations in expression of genes involved in regulating MuSC proliferation and differentiation, we quantified *c-Fos, Cdk4, MyoD*, and *Myog* mRNA expression. PFSS induced a 4-fold and 1.3-fold increase in *c-Fos* and *Cdk4* transcript levels, respectively (Fig. 4A). MuSCs expressed both *MyoD* and *Myog* with no difference in expression levels between PFSS-treated and static control cells (Fig. 4A). These results showed that PFSS induced expression of *c-Fos* and *Cdk4*, known to promote cell proliferation [25, 26].

**Figure 4.**
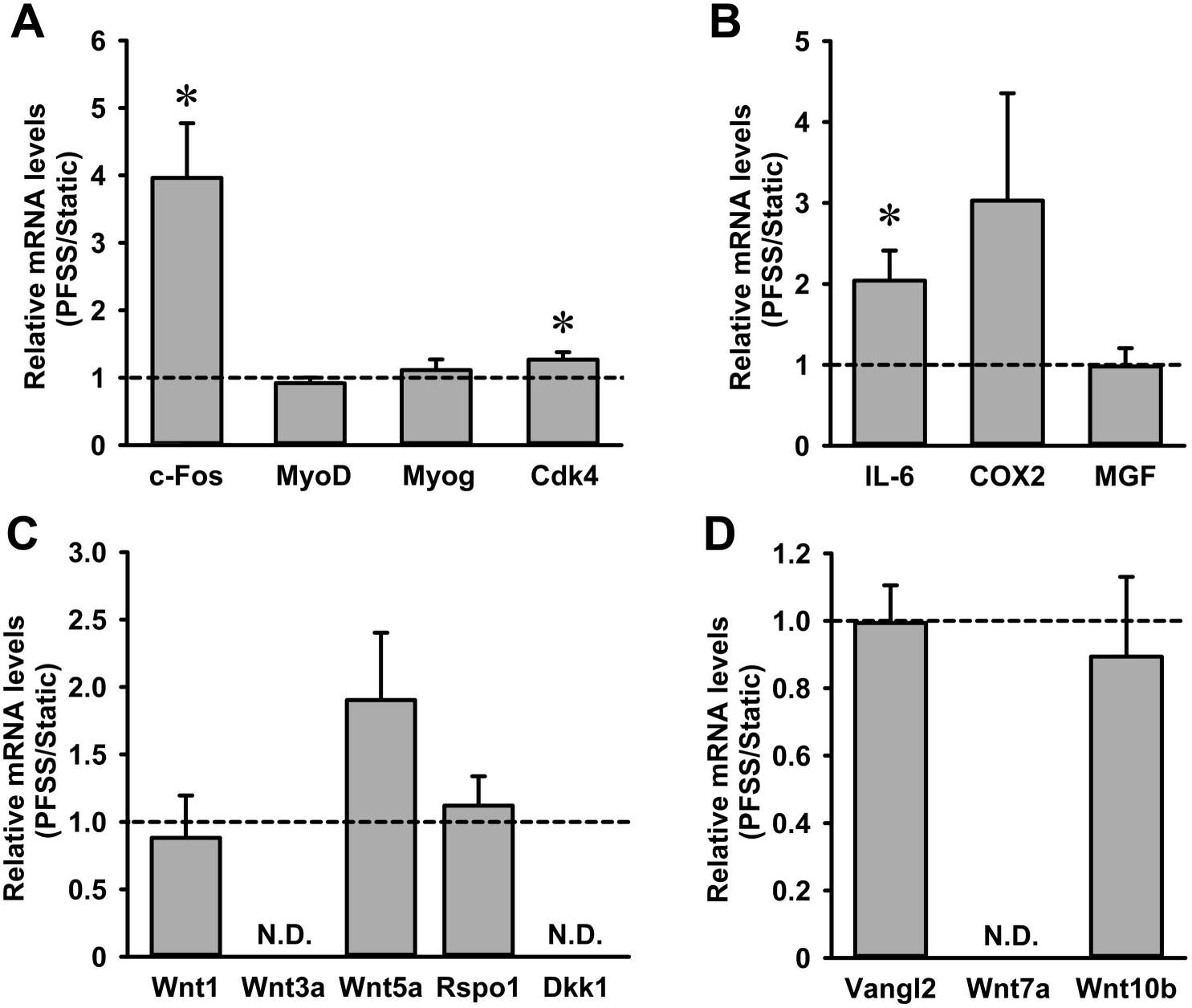
PFSS upregulated proliferation marker expression, i.e. *c-Fos, Cdk4*, and *IL-6*, in primary mouse MuSCs. (A) PFSS upregulated *c-Fos* expression by 4-fold, and *Cdk4* expression by 1.3-fold, but did not affect *MyoD* and *Myog* mRNA expression. (B) *IL-6* was upregulated by 2-fold after PFSS and an upward trend was observed in *COX2* expression, while no affect was shown on *MGF* by PFSS. (C) *Wnt5a* transcripts were slightly, but not significantly, upregulated, *Wnt1* and *Rspo1* expression remained unchanged, while *Wnt3a* and *Dkk1* were undetectable in both static and PFSS-treated cells. (D) PFSS did not affect *Vangl2* and *Wnt10b* expression, whereas *Wnt7a* was not detectable. n=7-10. PFSS, pulsating fluid shear stress; ND, not detectable. *Significant effect of PFSS, p<0.05.

### PFSS upregulated expression of *IL-6*, but not *COX2* and *MGF* in myoblasts

Exercise induces *IL-6* expression in myofibers, which is involved in myoblast proliferation and regeneration [27]. IL-6 production is stimulated by both NO and COX2-mediated prostaglandin production [28]. COX2 and mechano growth factor (MGF) are also essential for myoblast proliferation and differentiation [29, 30]. Therefore we determined the effect of PFSS on *IL-6, COX2*, and *MGF* expression in MuSCs. PFSS induced a 2-fold increase in *IL-6* expression and an upward trend towards increased *COX2* expression, but did not affect *MGF* expression (Fig. 4B).

### PFSS did not affect Wnt signaling

Wnt1 has been shown to stimulate MuSC proliferation [31]. To investigate the effect of PFSS on Wnt signaling within MuSCs, we evaluated the effect of PFSS on components of the Wnt signaling pathway (Wnt1, Wnt3a, and Wnt5a) and regulators of Wnt signaling, i.e. Rspo1 and Dickkopf-related protein 1 (Dkk1) [32]. PFSS did not affect *Wnt1* gene expression, while an upward trend was seen in *Wnt5a* mRNA levels (Fig. 4C). *Wnt3a* was undetectable, and *Rspo1*, a positive regulator of Wnt activity, was unchanged by PFSS, while *Dkk1*, a negative regulator, was undetectable (Fig. 4C). These results indicated that the Wnt signaling pathway was not affected by mechanical stimuli in MuSCs.

### PFSS did not alter expression of planar cell polarity genes

Wnt5a is implicated in mediating membrane protein Vangl2 phosphorylation and activation [33], known to affect the PCP pathway that is crucial for cell division and cell migration. We determined whether PFSS induced expression of *Vangl2* and showed that *Vangl2* expression was not affected by PFSS treatment (Fig. 4D). We further tested the effect of PFSS on *Wnt7a* and *Wnt10b* expression, which are involved in self-renewal and myogenic commitment of MuSCs. *Wnt7a* transcripts were not detectable in MuSCs, and *Wnt10b* expression was not affected by PFSS (Fig. 4D). Therefore, PFSS did not affect expression of genes involved in PCP formation in MuSCs.

## DISCUSSION

MuSCs within their native niche on the host myofiber with an intact endomysium, are presumed to undergo deformations when the myofibers are subjected to tensile strain. Here we have shown for the first time that MuSCs underwent both tensile strain and shear loading in response to myofiber strain. Due to constancy of myofiber volume, the endomysium surrounding the myofiber will undergo a reorientation from a longitudinal towards a more radial orientation upon muscle contraction [34]. Since MuSCs are anchored to the basal lamina and the sarcolemma of the host myofiber, one may expect that these cells also align in the orientation of basal lamina. Our data confirms such re-orientation of MuSCs with respect to the longitudinal axis of the myofibers.

Local differences in the stiffness of the endomysium may occur due to the presence of capillaries and differences in type of collagen and/or connective tissue content. These differences will cause shear loads and shear deformation of MuSCs upon mechanical loading of skeletal muscle. In both myofiber bundles of EDL and SO, we observed differences in lateral displacements along the length of MuSCs. This indicates that MuSCs are subjected to shear loading during myofibers stretch-shortening cycles. Mechanical loading of skeletal muscle may also cause localized changes in intramuscular pressure due to the movement of interstitial fluid, thus creating regions of different pressure gradients [35]. These pressure gradients potentially cause compression of MuSCs and result in altered cellular osmotic pressure and volume of MuSCs due to water efflux [36]. Osmotic pressure is a well-known regulator of mechanotransduction, implying that deformation of MuSCs by myofiber stretching induces activity in this manner. This study provides the proof of concept that MuSCs are subjected to forces and undergo different mechanical deformations which encourages further research on the nature of these forces in MuSC function and muscle regeneration.

Stretch of MuSCs results in the production of factors involved in MuSC activation, self-renewal, and proliferation [37]. We observed that shear-loading of MuSCs induced NO production and upregulated expression of *c-Fos, Cdk4*, and *IL-6*. NO is known to release hepatocyte growth factor (HGF) from the ECM and to mediate MuSC activation, proliferation, and fusion [37]. Since an upward trend in *iNOS* mRNA levels following PFSS application was observed, it is likely that *iNOS* is involved in PFSS-induced NO production in MuSCs. *c-Fos* is an early response gene and plays an important role in cell growth, proliferation, and differentiation [38]. During differentiation, c-Fos is repressed by MyoD and Myog [39, 40]. c-Fos is a subunit of transcription factor complex activator protein-1 (AP-1) [41], that binds to the *MyoD* promotor and inhibits *MyoD* transcription in proliferating myoblasts [42]. Overexpression of c-Fos has been shown to promote proliferation and inhibit differentiation in C2C12 myoblasts [43]. This suggests that increased *c-Fos* expression after shear-loading may induce MuSC proliferation. Moreover, CDK4 has been shown to play a role in cell cycle progression together with D-type cyclins [26]. IL-6 is essential for muscle regeneration and MuSC proliferation [27]. Muscle injury causes IL-6 production by macrophages and myofibers [4]. In this study, increased *IL-6* expression by shear-loaded MuSCs suggests a positive effect of loading on MuSC proliferation, and indicates that IL-6 may act in both a paracrine as well as an autocrine manner to induce MuSC proliferation. Moreover, expression of *COX2*, which is induced in response to injury and is involved in prostaglandin production and MuSC proliferation, also showed an upward trend after shear loading. Another important factor driving MuSC differentiation is MGF, which is induced in myotubes by PFSS [18]. We did not observe changes in *MGF* expression in MuSCs due to PFSS suggesting that mechanical loading for 1 h does not induce autocrine MGF signaling, whereas it does in differentiated myotubes [18]. This suggests that in addition to muscle damage and mechanical stretch, which stimulate MuSC proliferation, here we show that PFSS induced expression of proliferation-related genes and likely stimulates MuSC proliferation.

Shear-loading did not affect expression of genes associated with differentiation of myoblasts, as indicated by unaltered *MyoD* and *Myog* expression. Members of the Wnt family also govern MuSC function. Exogenous Wnt1 activates *Myf5* expression, resulting in MuSC activation and/or proliferation, as shown in rhabdomyosarcomas [44, 45]. Moreover, Wnt3a is required for both myogenic potential and differentiation of MuSCs [46]. We show that that *Wnt1* mRNA levels were unaltered in response to PFSS in MuSCs, whereas *Wnt3a* was not detectable in both controls and PFSS-treated cells. Wnt5a overexpression has also been shown to increase MuSC proliferation [47]. We show that MuSCs upregulate *Wnt5a* mRNA levels in response to PFSS, but the increase did not reach statistical significance. Wnt signaling may also be regulated by negative or positive mediators. Rspo1 mediates canonical Wnt signaling thereby affecting MuSC differentiation [48]. However, we did not observe changed *Rspo1* expression by PFSS. *Dkk1, a* negative regulator of Wnt signaling, was not detectable in MuSCs, suggesting that there was only a limited response by canonical Wnt signaling to PFSS, possibly involving upregulation of *Wnt5a*.

Wnt7a induces symmetric expansion of MuSCs by activating the PCP pathway through regulation of Vangl2, and is negatively regulated by Rspo1 [32, 49]. We did not detect *Wnt7a* expression in MuSCs, and *Vangl2* expression was unaffected by shear-loading, suggesting that PFSS did not affect PCP signaling in MuSCs. Wnt10b is involved in determining myogenic commitment of MuSCs as its ablation has been shown to decrease MuSC myogenicity [50]. PFSS did not alter *Wnt10b* expression suggesting no effect by PFSS on myogenic commitment of MuSCs.

## Conclusions

We conclude that MuSCs experience tensile strain, shear strain, and pressure in response to mechanical perturbations of the myofibers to which they are attached, and that mechanotransduction is likely to occur *in vivo* in MuSCs during muscle stretch-shortening. Shear-loading of MuSCs *in vitro* upregulated a number of genes that are known to promote MuSC proliferation (Fig. 5A,B). Fate mapping of MuSCs subjected to PFSS will further reveal whether this acts as a cue to induce MuSC proliferation and/or differentiation. Note that we observed that stretching of MuSCs also affected their cellular processes, thus potentially affecting their ability to migrate. Extended time-lapsed imaging of cells will therefore reveal how exposure to tensile and shear strain regulates MuSC proliferation, differentiation, and migration. This will provide important insight into how MuSCs respond to such forces in the context of ageing and disease, and how this affects their ability to maintain effective muscle function.

**Figure 5.**
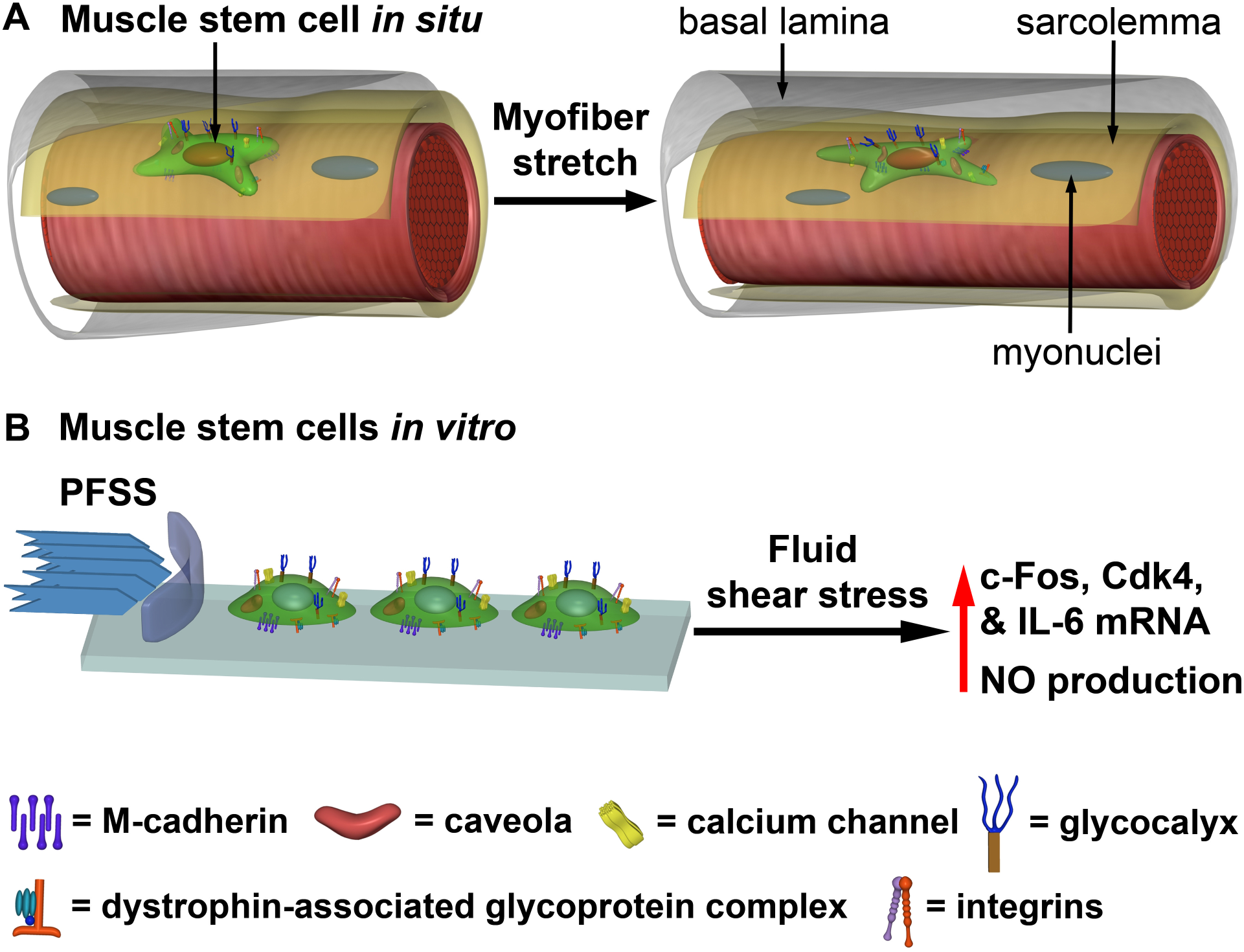
Schematic of myofiber stretch inducing MuSC deformation, and effect of fluid shear stress on myoblast mRNA expression. (A) MuSC in its niche on the host myofiber, embedded between the sarcolemma (yellow) and basal lamina (grey), and connected via transmembrane complexes, undergoing tensile strain and shear stress deformations upon myofiber stretch. (B) Myoblasts subjected to mechanical load by PFSS *in vitro* induce upregulation of NO and expression of factors known to promote proliferation.

## ABREVIATIONS

AP-1: Activator protein-1
Ca^2+^: Calcium
Cdk4: Cyclin-dependent kinase 4
COX2: Cyclooxygenase-2
Dkk1: Dickkopf-related protein 1
ECM: Extracellular matrix
EDL: Extensor digitorum longus
eNOS: Endothelial nitric oxide synthase
FBS: Fetal bovine serum
HGF: Hepatocyte growth factor
IL-6: Interleukin-6
iNOS: Inducible nitric oxide synthase
mG: membrane targeted green fluorescent protein
MGF: Mechano growth factor
MITK: Medical Imaging Interaction Toolkit
mT: Membrane-targeted tandem dimer Tomato
MuSCs: Muscle stem cells
MyoD: Myogenic differenciation
Myog: Myogenin
nNOS: Neuronal nitric oxide synthase
NO: Nitric oxide
NO_2_ ^−^: Nitrite
NOS: nitric oxide synthase
Pax7: Paired box 7
PCP: Planar cell polarity
PFSS: Pulsating fluid shear stress
Rspo1: R-spondin-1
SO: Soleus
Vangl2: Vang-like protein 2

## DECLARATIONS

### Ethics approval and consent to participate

Animal procedures were conducted according to the UK Home Office project license P8D5E2773 (KJL) and the guidelines of the European Community and French Veterinary Department, approved by the Sorbonne Université Ethical Committee for Animal Experimentation.

### Consent for publication

Not applicable.

### Availability of data and materials

All data generated or analysed during this study are included in this published article.

### Author contributions

MH, JJ, KJL, RDK, and RTJ performed the experiments. MH, ADB, CO, and RDK analyzed the data. MH, JJ, and RTJ drafted the manuscript. MH, JKN, ADB, JJ, CO, FLG, LG, KJL, RDK, and RTJ contributed to conceive and plan the experiments, interpret the results, and edited and revised the manuscript.

## Acknowledgements

The authors thank Dylan Herzog for technical assistance.

## ADDITIONAL FILES

### Additional file 1

**Figure S1**. Chamber for incubation of myofiber bundle at slack length or high strain, consisting of a glass bottom and adjustable platinum rods.

File name: Haroon et al Additional file 1

File format: Microsoft Word Document (.docx)

Title of data: Supplemental Information

Description of data: Contains supplementary Figure S1 legend.

