## Supplementary figures and images for "Myofiber stretch induces tensile and shear deformation of muscle stem cells in their native niche"

### Figure S1

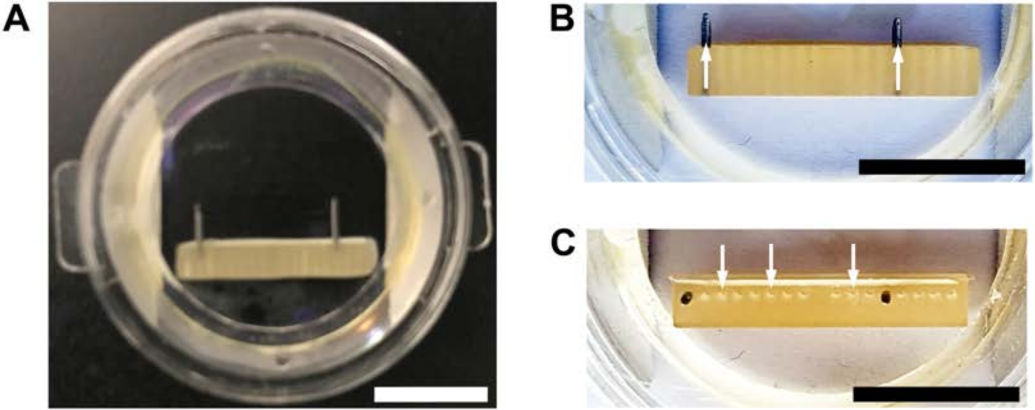
