## Supplemental Information for "Myofiber stretch induces tensile and shear deformation of muscle stem cells in their native niche"

**Figure S1.** Chamber for incubation of myofiber bundle at slack length or high strain, consisting of a glass bottom and adjustable platinum rods. (A) Chamber used to maintain the myofiber bundle at slack length or high strain. (B) The platinum rods (arrows) in the chamber used to mount the myofiber bundles. (C) The inserts (arrows) in the bar holder allow to maintain the myofiber bundle at slack length or high strain by adjusting the distance between the rods. Scale bar, 1 cm.
